# The similarities in microbial and chemical patterns of fermentation in two open environments were promoted by using 150-year-old Nukadoko as starters

**DOI:** 10.1101/2023.11.17.567490

**Authors:** Marin Yamaguchi, Kota Imai, Dominique Chen, Young ah Seong, Kazuhiro Jo, Kohei Ito

## Abstract

Nukadoko, a fermented rice bran employed in traditional Japanese pickling, uses lactic acid bacteria to ferment vegetables. Here, we report the microbial and chemical data of a mixture of matured 150-year-old nukadoko and commercially available rice bran placed in two open environments over 29 days. Across the two environments, *Loigolactobacillus* was identified as the dominant microbial genera in the later stages of fermentation in nukadoko. The period of increase in the relative abundance of *Loigolactobacillus* correlated with a decrease in pH and Oxidation-Reduction Potential (ORP) values. While the two environments showed a difference in the rate of change in microbial diversity, they shared the common process through which *Loigolactobacillus* outcompeted adventitious bacteria in nukadoko, as indicated by the alpha and beta diversity index. Thus, the similarities in microbial and chemical data across two open environments during fermentation using starters indicate that starters contribute to the stability of fermentation in open environments.

## Introduction

Nukadoko, a fermented rice bran employed in traditional Japanese pickling, uses lactic acid bacteria (LAB) to ferment vegetables (*Nukazuke*). Traditionally, the nukadoko preparation process involves the natural fermentation of rice bran, continuously renewed by adding fresh bran. A previous study reported that the initial fermentation time was thereby reduced using an aged nukadoko as a fermentation starter ready for use (Ono et al., 2014). Indeed, many modern fermentation practices often involve the use of starters, which also serve the purpose of preventing food spoilage in the fermentation process of meat, fish, and dairy products (2).

The microbiota of the fresh rice bran used in nukadoko is especially rich in microbial diversity, giving rise to the distinct flavor of nukadoko (Ono et al., 2014). The type of rice bran used, salinity levels, temperature, air exposure, and fermentation time can all be factors that distinguish the nukadoko microbiota (3). However, in a previous study involving four types of nukadoko, such microbial diversity eventually converges to represent similar microbial communities as the fermentation process continues (Ono et al., 2014). A similar phenomenon has also been observed in other fermented foods such as kimchi, where samples of various types of kimchi at later stages of fermentation were mainly composed of *Lactobacillales*, related to LAB (4). However, still little is known about the complete picture of the nukadoko microbiota, as well as the chemical changes during the fermentation process.

In this study, we examined the chemical and microbial changes during the fermentation of nukadoko in two different open environments over 29 days.

## Materials and Methods

### Preparation of *nukadoko* and sample collection

Nukadoko utilized in this study was prepared by mixing commercially available rice bran (Maruho Foods, Kagawa, Japan) with matured 150-year-old nukadoko. The long-aged (approximately 150 years) nukadoko used in this experiment were obtained from a nukadoko manufacturer (Hyaku-Gojunen-no-Nukadoko-Hozonkai) in Fukuoka City, Japan. The ingredients of the 150-year-old nukadoko were rice bran, salt, water, *kombu* (a type of edible seaweed), dried young sardines, bird’s eye chili, Sichuan peppercorns, and rice malt. 250 g of the 150-year-old nukadoko was mixed with fresh rice bran (500 g), salts (125 g), and distilled water (500 mL). The mixed samples were placed in two locations for 29 days.

Three participants at both Location 1 and Location 2 were assigned to stir nukadoko on a daily basis at each location (Fig. 1). We chose Location 1, Hakko Department, a commercial establishment specializing in fermented foods and also offering dining options, and location 2, an exhibition at The National Museum of Emerging Science and Innovation, as the experimental locations. We deployed two sets of nukadoko vessel equipped with pH and ORP sensors (27) to each location. Among Location 2, a public space where temperature and humidity are strictly controlled, and Location 1, a more enclosed environment, we aimed to investigate whether the patterns of nukadoko fermentation vary among the two locations with such different characteristics and user demographics. The frequency of stirring was determined according to the hygiene standards of each facility and the operating patterns of the staff: once daily at Location 1 and twice daily at Location 2, once every morning and once every evening. For each stirring session, the participants were asked to mix nukadoko thoroughly from the bottom 2-3 times, with each session spanning approximately 3 minutes.

**Fig. 1.**
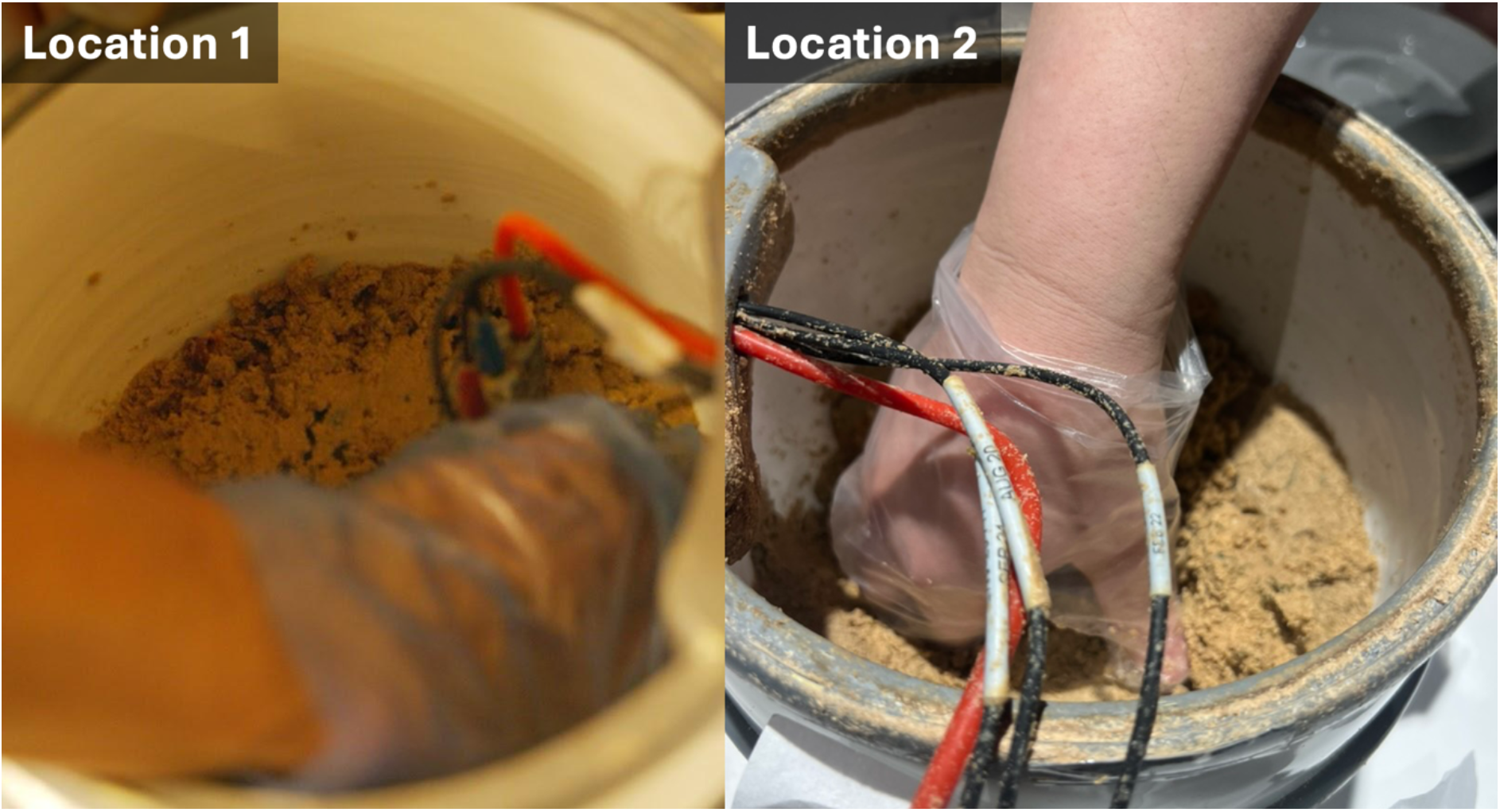
Pictures demonstrating the maintenance of nukadoko in two open environments.

Sample collection took place on specific days, namely days 0, 3, 6, 9, 15, 17, 22, and 29 for Location 1, and on days 0, 3, 6, 9, 14, 18, 22, and 29 for Location 2, using individually wrapped disposable plastic spoons and 50mL polypropylene conical tubes. Thus, we obtained 16 samples in total from the two locations. Samples were stored in a freezer after collection and kept at -20°C until DNA extraction.

### Chemical analysis

We employed high-precision pH and ORP sensors for robots developed by Atlas Scientific (USA) and inserted them into the nukadoko rice bran. These sensors were connected to electronic circuits (Raspberry Pi 4) that transmit the sensed data to a cloud database (Amazon Web Service RDS) each minute during the whole period.

### Total DNA extraction and high-throughput sequencing

DNA isolation from 50 ml nukadoko samples was performed using the beat-beating method using the multi-beads shocker (Yasui Kikai, Osaka). The extracted DNA was quantified using Qubit™ (Thermo Fisher Scientific, Massachusetts). The DNA was further purified using a GenCheck? DNA Extraction Kit Type S/F. The first PCR cycle was conducted using ExTaq HS DNA Polymerase, where specific primers 341F (5’-CCTACGGGNGGCWGCAG-3’) and 805R (5’-GACTACHVGGGTATCTAATCC-3’) of the V3-V4 region of the 16S DNA were amplified (5). 2 μl of the first PCR product, purified using AMPure XP beads, served as a template for library preparation. The second PCR product was again purified using AMPure XP beads. Amplicon sequencing was performed using 460 bp paired-end sequencing on the Illumina MiSeq platform (Illumina Inc., San Diego, CA, USA) at FASMAC Co., Ltd (Kanagawa, Japan).

### Microbial analysis

Microbial analysis was performed as previously regarding previous studies (6). Briefly, raw FASTQ data were imported into the QIIME2 platform (2022.8) as qza files (7). Denoising sequences and quality control were performed using the QIIME dada2, and finally, sequences were produced into amplicon sequence variants (ASVs) (8). ASVs were assigned to the SILVA database’s SSU 138 using the QIIME feature-classifier classification scikit-learn package (9,10). ASVs classified as mitochondria, chloroplast, or unassigned were excluded from subsequent analysis. Subsampling was performed to reduce bias due to differences in read depth between samples. The beta diversity indices weighted and unweighted UniFrac distances were calculated, and the microbial community structure differences between the two groups were visualized following a principal coordinate analysis (PCoA). Data were visualized using R (version 4.0.4), ggplot2 (version 3.4.3), and ggprism (version 1.0.4) (11,12).

### Statistical analysis

Mann–Whitney U tests were used to compare all combinations of elements within the two locations.

### Data Availability

The datasets generated through 16S rRNA amplicon sequencing are available and deposited in the DDBJ Sequence Read Archive (SRA) database under accession numbers DRR514074-DRR514089 and BioProject PRJDB17051.

## Results

### Changes in ORP and pH values during fermentation of nukadoko

Fig. 2a shows the gradual decrease in pH values for both samples, indicating the fermentation rate. The most rapid decrease in pH occurred between days 14 and 16 for Location 1 and between days 3 and 7 for Location 2 (Fig. 2a). The pH values indicated the difference in the time period in which rapid fermentation occurs among the two sample locations. Finally, the pH values at the two locations on the last day (day 29) were significantly different (p < 0.05).

**Fig. 2.**
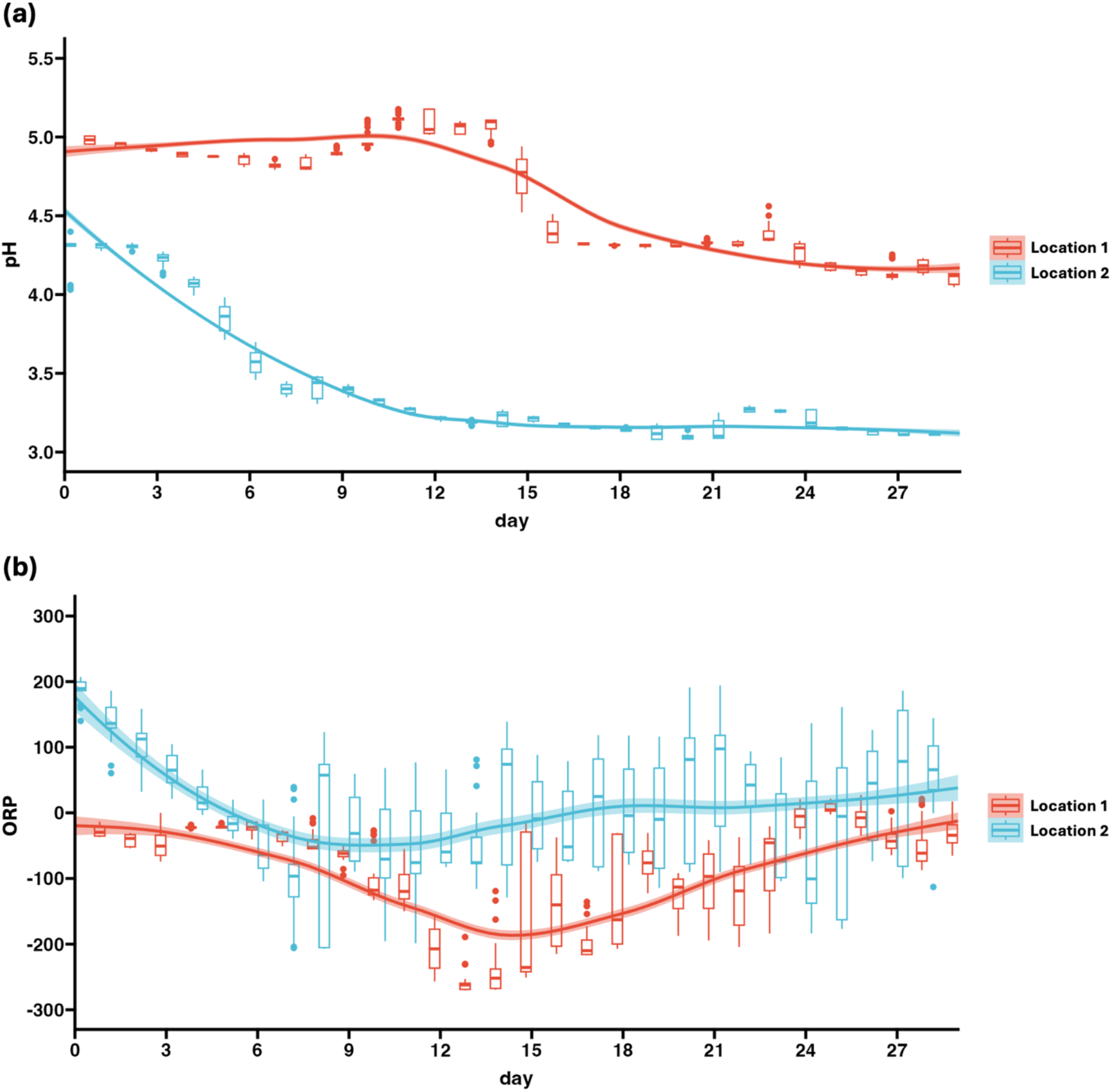
Regression plots of time-series data of OPR (a) and pH (b) values of the nukadoko environment in Location 1 and Location 2 samples over 29 days. The red line represents data from the Location 1 sample, and the blue line represents data from the Location 2 sample.

The ORP values showed variability throughout the experiment. The decline in the ORP values from approximately -50 mV to -250 mV occurs in the Location 1 sample across 5 days (days 9-13), and the decline from approximately 200 mV to -100 mV occurs in the Location 2 sample across 8 days (days 0-7) (Fig. 2b). Similarly to the results obtained through the differences in pH values, the variation in ORP across the experiment is also indicative of the different nature of the fermentation process among Location 1 and Location 2. Finally, the ORP values at the two locations on the last day (day 29) were not significantly different (*p* = 0.0507).

### Increase of the relative abundance of LAB

The number of raw reads obtained from 16 samples remaining after quality filtering was 323,217. After removing ASVs classified as mitochondrial, chloroplast, or unassigned, the total number of reads was 201,452. The ASVs from the samples were classified, with the most abundant genera being *Loigolactobacillus, Pantoea, Sphingomonas, Allorhizobium-Neorhizobium-Pararhizobium-Rhizobium, Pseudomonas*, and *Staphylococcus* on day 0 (Fig. 3). Over time, *Loigolactobacillus* dominates the majority in both Location 1 (Day 0: 69.6 % and Day 29: 99.0 %) and Location 2 (Day0: 69.5 % and Day 29: 99.4 %) samples. Despite the similar pattern, the rate of decrease in microbial diversity follows different paths, as the Location 1 sample continued to exhibit a wider variety of microbes over a longer period, whereas, in the Location 2 sample, *Loigolactobacillus* dominated the majority at a much earlier phase, at around day 6 of the experiment.

**Fig. 3.**
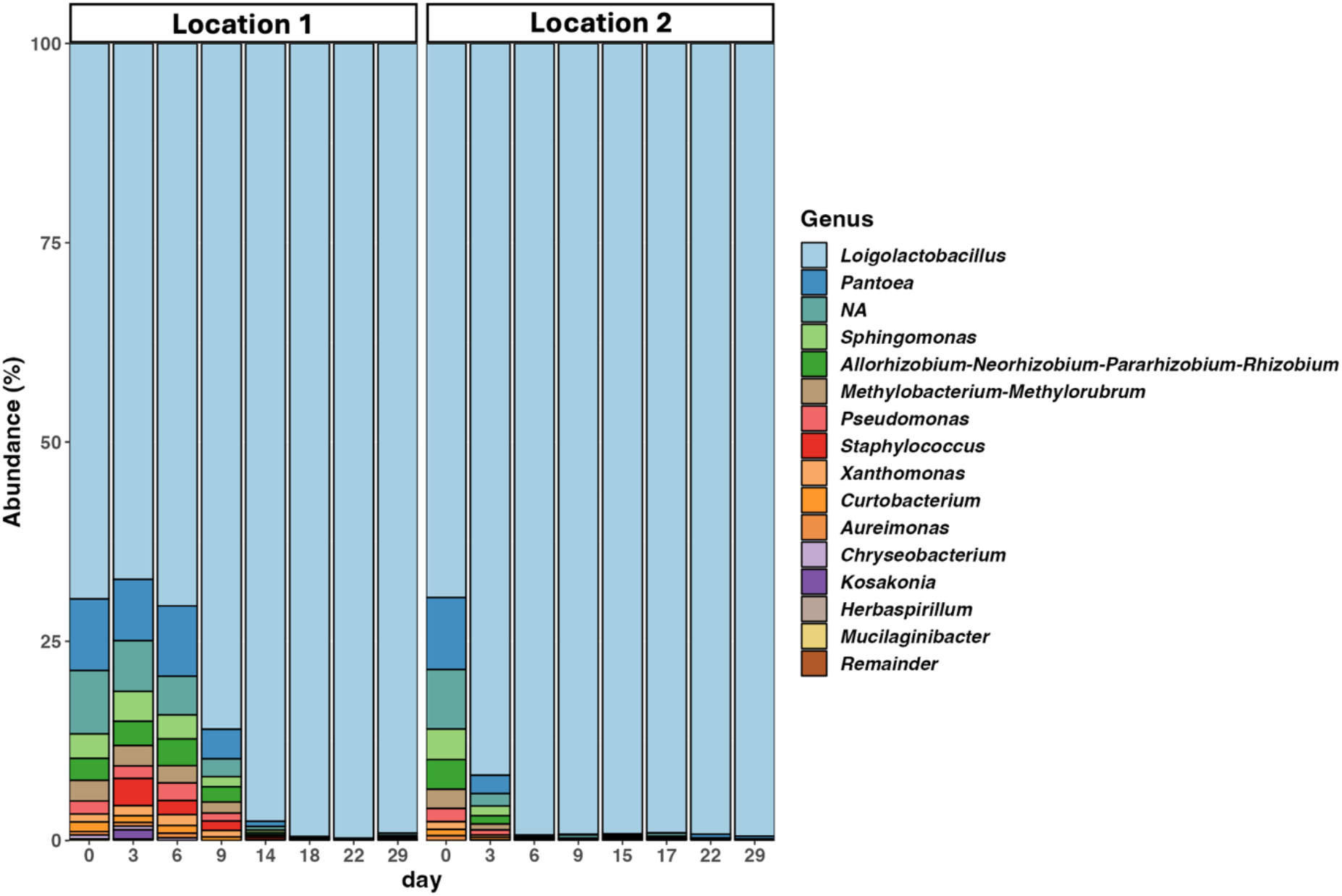
Stacked bar plots of the genus-level taxonomic composition during the fermentation of nukadoko over 29 days. The top 15 genera are listed, and the rest are noted as Remainder.

The microbial community structures of both samples were compared using principal component analysis (PCoA) by weighted UniFrac distance metrics. At the beginning, both Location 1 and Location 2 show similar microbial compositions, consisting of a broad range of microbial genera, as shown in Fig. 3. This is indicated by the proximity with which the two sample points on day 0 are located on the PCoA plot. However, between days 3 and 9, Location 1 and Location 2 samples exhibited distinct microbial compositions and eventually showed similarities again towards the end of the experiment (Fig. 4a). Similarly to the results indicated by Fig. 2a, this indicates the difference in the process and rate of fermentation among the two sample locations. Indeed, at around day 3, the sample point from the Location 2 sample approaches the clustered sample groups, indicating the end of the fermentation process. However, in the Location 1 sample, the data points started to cluster together on day 14, displaying the gap in fermentation rate among the two sample locations. The sample points at the end of fermentation can be clustered together, where PC2 measures around 0%, indicating that the fermentation process exhibit similar microbial trends across the two locations over time. The Shannon Entropy values also exhibit similar results, where the decrease in the alpha diversity occurs exponentially, and then eventually stabilizes. The alpha diversity measures high among the Location 1 sample until day 14 (Fig. 4b), and its decrease at the end coincides with the period *Loigolactobacillus* takes over as the most dominant genus. The days in which the alpha diversity decreases correlate with the days *Loigolactobacillus* dominates, occurring between days 14-15 for the Location 1 sample and between days 6-7 for the Location 2 sample.

**Fig. 4.**
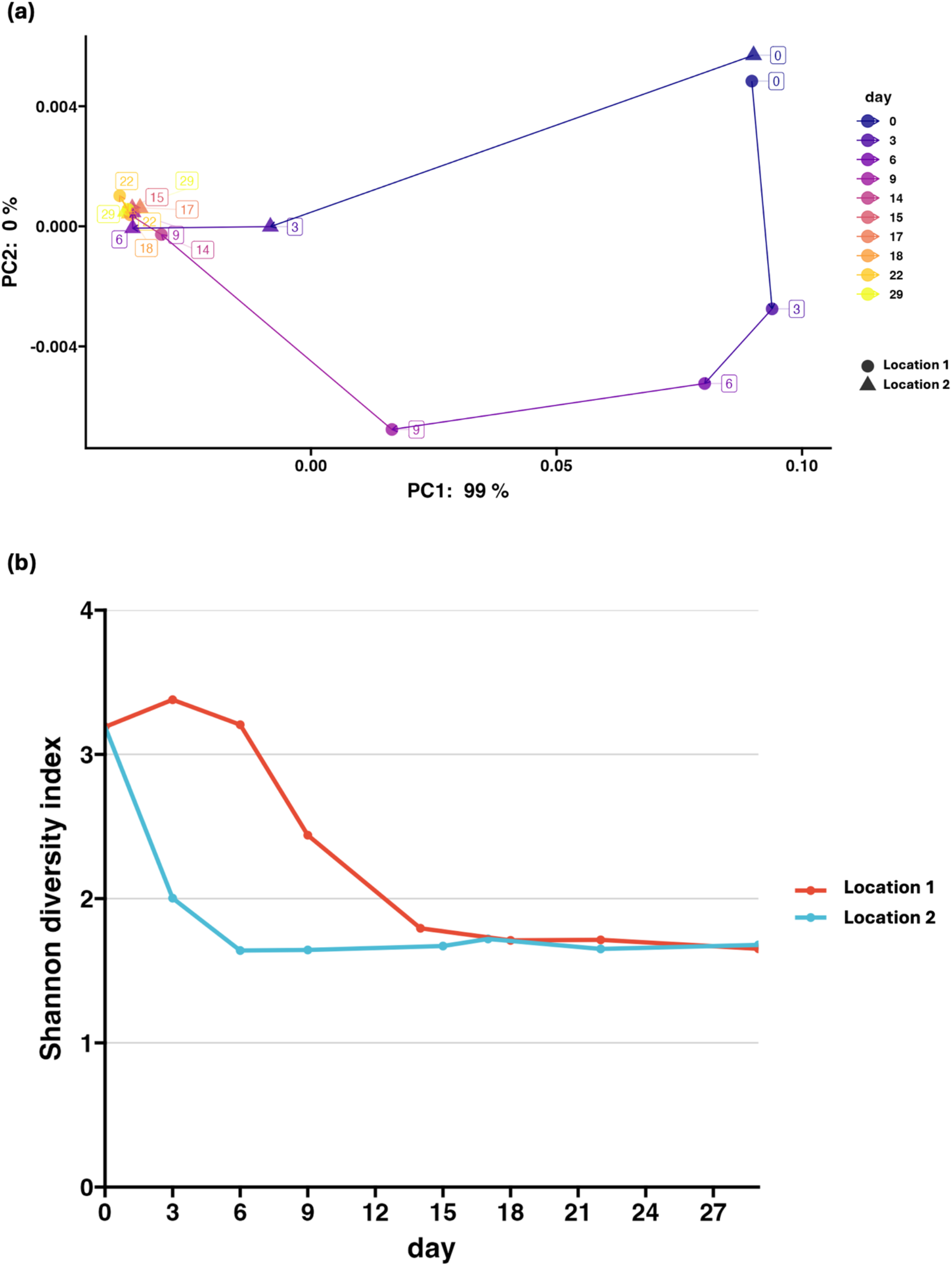
Changes in the microbial diversity during the fermentation of nukadoko samples. (a) Principal-coordinate analysis (PCoA) by Weighted UniFrac distance metrics of Location 1 and Location 2 samples. The line connects the data points across 29 experimental days. The data points represented by the circles indicate Location 1, and data points represented by the triangles indicate Location 2. (b) Line plots of Shannon diversity index of Location 1 and Location 2 samples over 29 days.

This period of change in microbial diversity aligns with the period of changes in pH values obtained through chemical analysis.

## Discussion

### Changes in the ORP and the pH values during fermentation of nukadoko

The ORP values are often used to evaluate the micro-aerobic status of the environments in which fermentation take place (13). In a previous study involving the *Lactobacillus mucosae* species the ORP values of micro-aerobic fermentation was determined to be between -300 and -150 mV (14). The readings of ORP values obtained in this experiment mainly lie within such micro-aerobic fermentation level, indicating the normal fermentation process induced by the daily mixings and the concurrent air intake into nukadoko. This is also consistent with previous findings that ORP values reflect activities of both aerobic (positive ORP) and anaerobic (negative ORP) bacteria, and thus ORP values are indicative of the level of fermentation (15). While aerobic growth may promote oxidative stress in LAB species, oxygen availability is an important factor that determines the level of aromatic compounds produced by LAB (16). Aromatic compounds, mainly acetoin and diacetyl, are known to impact the flavor of dairy products during food fermentations with LAB (17), suggesting that the distinct flavors of nukadoko may also be supported by similar aromatic compounds. Thus, the micro-aerobic environments in the two locations in this experiment may have influenced the metabolic routes of LAB to produce acetoin and diacetyl, giving rise to the nukadoko flavor.

A previous study has also demonstrated the abundance of LAB and the accompanying pH decrease during the late fermentation period, similar to the results obtained in this experiment (18). Among the two sample locations, the time points of chemical and microbial changes are aligned, demonstrating the advancement of nukadoko fermentation. Changes in chemical values including pH values occurred between days 14-16 in Location 1 and days 3-7 in Location 2, which coincide with the days in which alpha diversity decreases for both locations. This suggests that the chemical and microbiota data of nukadoko fermentation are interrelated.

### Changes in the taxonomic composition during fermentation of nukadoko

Our results showed that *Loigolactobacillus*, an anaerobic, homo-fermentative bacteria, became dominant in both environments. Similarly, in another study, *Loigolactobacillus* was identified as the dominant genus in nukadoko samples (19). The process through which adventitious bacteria are outcompeted by LAB at the later stages of fermentation is dependent on the types of LAB that dominate the microbiota (17). A previous study (18) investigated the chemical changes during the initial stage of fermentation across four samples made of different types of rice bran. Lactate concentration can be attributed to the differences between hetero-fermentative LAB during the early fermentation stage and homo-fermentative LAB at the later stages (19). The dominance of *Loigolactobacillus* in our experiment suggests the abundance of homo-fermentative bacteria in 150-year-old nukadoko, indicating its matured fermentation state. It must be noted that *Loigolactobacillus* is a genus deriving from the reclassification of *Lactobacillus* (20).

In another study, Sakamoto identified two predominant species of *Lactobacillus* that maintain a nukadoko environment that prevents contamination by other microorganisms, conserving the distinct nukadoko flavor and preventing further spontaneous fermentation by adventitious microbes (21). Indeed, within the architectural environment in which the samples are placed, various microbes, known as the Microbiome of the Built Environment (MoBE), exist, which can influence the production of fermented foods (22,23). MoBE has been found to contaminate the fermentation starter in a *sake* brewery (24). Similarly to the effects of *Lactobacillus* dominance of the microbiota in other experiments involving nukadoko, the presence of *Loigolactobacillus* in our experiment may also provide stability to the fermentation process and prevent adventitious microbes from contaminating the nukadoko environment. The high prevalence of *Loigolactobacillus* from day 0 may have resulted from an increased amount of readily matured nukadoko. We used an increased amount of starter compared to a previous study (18) in order to support fermentation in open environments.

Bacterial genus recently isolated from nukadoko in other findings, including Weisella (doi: 10.1128/MRA.01160-21) and Bacillus (doi: 10.1128/MRA.00705-21), were rarely present in the nukadoko samples. This suggests the wide variety of microbiota in different nukadoko environments.

We also examined the microbial composition of nukadoko over the fermentation period, where most adventitious bacteria such as *Pantoea, Pseudomonas*, and *Staphylococcus* originating from sufficiently aged nukadoko were not detected by the end of the fermentation process in both Location 1 and Location 2 samples. Such changes in microbial composition resemble data obtained through previous studies, where one (24) notes the changes from the prevalence of *Enterobacter*, a type of gram-negative bacteria encompassing *Pantoea*, to that of LAB in the later stages of fermentation. Thus, adventitious bacteria are continuously outcompeted by *Lactobacillus* and/or *Loigolactobacillus* and their lactic acid products. The LAB-dominant microbiota is not unique to nukadoko and can be seen in several fermented foods (25).

### The microbial diversity of the two locations eventually became similar

Fig. 4a indicates that the two groups exhibit differences in the sample microbiota at the initial stages of fermentation but later show high similarities. Fig. 4b indicates the gradual decrease in the Shannon Diversity index across two groups at the later stages of fermentation. It is suggested that these changes in microbial diversity reflect the gradual dominance of *Loigolactobacillus* throughout the fermentation process. This is consistent with the findings in other fermented foods, including Suancai, a traditional sauerkraut in Northeast China. In an experiment involving Suancai fermentation, 16S rRNA microbial analysis indicated that by day 30 of fermentation, 100% of the sample bacteria comprised of LAB (26). Indeed, the dominance of LAB at later stages of fermentation is often observed in various types of fermented foods, which reflects the common modern fermentation practice of introducing starters so that fermentation-contributing microbes can dominate the microbiota at a more rapid pace (2).

It must be noted that *Loigolactobacillus* was detected at a relatively high abundance in the nukadoko from the initial stage of fermentation, which may have been caused by an excessive amount of starter. In addition, due to the analysis at the ASV levels, it is yet to be known whether the detected *Loigolactobacillus* consists of a single or multiple species.

## Conclusion

Through microbial and chemical analyses, we have demonstrated the similarities in microbial and chemical patterns during nukadoko fermentation in two different open environments. Such similarities in the fermentation process observed among the two sample locations demonstrates how the 150-year-old nukadoko starter contributes to the creation of the optimal environment for nukadoko fermentation, characterized by the dominance of *Loigolactobacillus*.

## Acknowledgements

We would like to thank “Hyaku-goju-nen no Nukadoko Hozonkai” for providing their matured Nukadoko and giving us permission to analyze its microbiota.

## Statements & Declarations

### Author Contributions

The study was conceived by all the authors. Chen Dominique and Kohei Ito designed the experiments. Chen Dominique collected samples. Marin Yamaguchi drafted the original manuscript. Kota Imai and Kohei Ito performed microbiome analysis. Young ah Seong, Kazuhiro Jo, and Chen Dominique edited the manuscript and supervised the study. All authors have contributed to the manuscript and approved the submitted version.

### Ethics Approval and Consent to Participate

The research protocol was approved by the Research Ethics Committee, Faculty of Design Engineering, Hosei University (Application Number:2022-4; approval date: March 28, 2023). All procedures were conducted according to the ethics committee’s guidelines and regulations. Informed consent was obtained from all participants before starting this study.

